# Heterologous booster with a novel formulation containing glycosylated trimeric S protein is effective against Omicron

**DOI:** 10.1101/2023.10.05.557343

**Authors:** Daniela Bottero, Erika Rudi, Pablo Martin Aispuro, Eugenia Zurita, Emilia Gaillard, Maria M. Gonzalez Lopez Ledesma, Juan Malito, Matthew Stuible, Nicolas Ambrosis, Yves Durocher, Andrea V. Gamarnik, Andrés Wigdorovitz, Daniela Hozbor

## Abstract

In this study, we evaluated the efficacy of a heterologous three-dose vaccination schedule against the Omicron BA.1 SARS-CoV-2 variant infection using a mouse intranasal challenge model. The vaccination schedules tested in this study consisted of a primary series of 2 doses covered by two commercial vaccines: an mRNA-based vaccine (mRNA1273) or a non-replicative vector-based vaccine (AZD1222/ChAdOx1, hereafter referred to as AZD1222). These were followed by a heterologous booster dose using one of the two vaccine candidates previously designed by us: one containing the glycosylated and trimeric spike protein (S) from the ancestral virus (SW-Vac 2μg), and the other from the Delta variant of SARS-CoV-2 (SD-Vac 2μg), both formulated with Alhydrogel as an adjuvant. For comparison purposes, homologous three-dose schedules of the commercial vaccines were used. The mRNA-based vaccine, whether used in heterologous or homologous schedules, demonstrated the best performance, significantly increasing both humoral and cellular immune responses. In contrast, for the schedules that included the AZD1222 vaccine as the primary series, the heterologous schemes showed superior immunological outcomes compared to the homologous 3-dose AZD1222 regimen. For these schemes no differences were observed in the immune response obtained when SW-Vac 2μg or SD-Vac 2μg were used as a booster dose. Neutralizing antibody levels against Omicron BA.1 were low, especially for the schedules using AZD1222. However, a robust Th1 profile, known to be crucial for protection, was observed, particularly for the heterologous schemes that included AZD1222. All the tested schedules were capable of inducing populations of CD4 T effector, memory, and follicular helper T lymphocytes. It is important to highlight that all the evaluated schedules demonstrated a satisfactory safety profile and induced multiple immunological markers of protection. Although the levels of these markers were different among the tested schedules, they appear to complement each other in conferring protection against intranasal challenge with Omicron BA.1 in K18-hACE2 mice. In summary, the results highlight the potential of using the S protein (either ancestral Wuhan or Delta variant)-based vaccine formulation as heterologous boosters in the management of COVID-19, particularly for certain commercial vaccines currently in use.

## Introduction

Severe acute respiratory syndrome coronavirus-2 (SARS-CoV-2) is the virus that causes the pandemic coronavirus disease 2019 (COVID-19) with over 768 million infections and 6.9 million deaths to date (https://covid19.who.int/). Unprecedented in the history of vaccinology, within less than 12 months from the start of the pandemic, several research teams successfully developed vaccines that generated protective immunity against COVID-19 (1–4). Indeed, it was primarily the vaccines that altered the trajectory of the pandemic, significantly decreasing not only the incidence of severe cases but also the fatalities associated to SARS-CoV-2 infection. The first COVID-19 vaccine was administered outside of a clinical trial setting on December 8, 2020. During the first year of COVID-19 vaccine distribution, approximately 56% of the global population has received at least one dose of a COVID-19 vaccine, with 45.5% estimated to have received two doses, and 4.3% estimated to have received a booster (5). Based on official reported COVID-19 deaths, it was estimated that vaccinations prevented 14.4 million deaths from COVID-19 in 185 countries and territories between December 8, 2020, and December 8, 2021 (5). However, the pandemic still continues and there are several challenges to overcome. One of them is to ensure equal availability of these vaccines for people all over the world. Although more than 13.5 billion doses have already been administered worldwide, only 32.2% of people in low-income countries have received at least one dose (https://ourworldindata.org/covid-vaccinations). Another challenge refers to the evolution of SARS-CoV-2 and its impact in the public health. Since the beginning of the COVID-19 pandemic, the World Health Organization (WHO) has designated multiple variants of SARS-CoV-2 as variants of concern (VOCs) based on the assessment of their transmissibility, ability to replace previous variants and cause new increases in infections with higher circulation, as well as the need to adjust public health measures in terms of prevention. Until now, there were 5 SARS-CoV-2 VOCs as defined by the WHO: alpha (pangolineage: B.1.1.7), Beta (B.1.351), Gamma (P1), Delta (B.1.617.2) and Omicron (B.1.1.529 and descendant lineages). These variants have mutations with functional significance in the gene responsible for expressing the glycoprotein spike. The significance of the accumulation of mutations in the RBD region of the spike protein is due to the role that this region plays in the main process of virus entrance into host cells via the angiotensin-converting enzyme 2 receptor (ACE2) and because it is the main target of neutralizing antibodies (6,7).

Based on comparisons of antigenic cross-reactivity using animal sera, replication studies in experimental models of human respiratory pathways, and clinical and epidemiological studies in humans, there is a consensus among experts in the Technical Advisory Group on the Evolution of the SARS-CoV-2 Virus that the Omicron variant represents the most divergent variant seen to date, when compared to previous variants (8). Since its appearance, Omicron has continued to evolve and has led to an increasing range of sublineages, all of which have been characterized so far by their ability to evade existing population immunity and to preferentially infect the upper respiratory tract (as opposed to the lower). Since Omicron variant and its different sublineages became dominant, the main focus of the vaccine strategy has been the enhancement of neutralizing antibodies (NAb) through vaccines booster doses. This is because NAbs are considered an important correlate of protection (9–11). It was also observed that hybrid immunity (natural infection plus vaccination) induces higher NAb levels than vaccine-only immunity (12). While the difference in immune response between receiving a booster or not is clear, there is currently a concern regarding the concept of immunological original sin (13). Recent studies have demonstrated how immunological imprinting shapes individuals’ responses to SARS-CoV-2. It has been observed that individuals infected with the earliest strain or subsequent Alpha or Beta strains exhibit varying immune responses to a subsequent Omicron infection, depending on their initial exposure. Moreover, exposure to the Omicron strain itself does not appear to update the imprinted response in individuals previously infected or vaccinated with an older strain, which may explain reinfection. In line with these findings, the study published by Alsouss et al. (14) detected a high percentage of circulating memory B cells (MBCs) that recognize the S protein from the ancestral SARS-CoV-2 variant in individuals boosted with variant S antigens. While immunization with a distinct antigen can activate rare naive B cells that target novel epitopes, the overall response continues to be dominated by clones that cross-react with the original strain. This means that MBCs generated during the primary vaccination series dominate the recall response induced by the booster, potentially outcompeting clones specific to novel epitopes. The way to counteract the effects of original sin is not clearly defined. One potential approach could involve administering multiple booster doses of monovalent vaccines containing immunogens from prevalent circulating variants. However, the availability of such vaccines may be delayed due to the need to determine the prevalence of these variants, especially over time, and to assess their impact on immunogenicity. While these developments are underway, alternative options could involve broader platforms in terms of epitopes and/or heterologous regimens using already existing vaccines.

In this study, we present findings on the protective effect against Omicron variant and immunogenic characterization of three dose heterologous vaccination protocols. The schedules involved a booster dose containing a vaccine candidate based on the soluble trimeric prefusion stabilized spike domain of the ancestral or Delta variants. The booster was applied to a primary vaccination series of 2 doses of commercially available vaccines.

## Materials and Methods

### Recombinant S protein from different SARS-CoV-2 variants

Recombinant Spike trimer constructs are based on “tagless” versions of the SARS-CoV-2 Spike trimers described previously (15,16). Briefly, SARS-CoV-2 sequences (Genbank accession number MN908947) were codon-optimized for Chinese Hamster Ovary (CHO) cells and synthesized by GenScript. Within the construct, the spike glycoprotein was preceded by its natural N-terminal signal peptide and fused at the C-terminus to the foldon domain of the T4 phage head fibritin (17). Mutations were added to stabilize the generated spike protein in the pre-fusion conformation (K986P–V987P) and the furin site was abolished (RRAR^682–685^ –GGAS) as previously described (18). Expression of the pTT241 plasmid constructs that did not encode C-terminal FLAG/His affinity tags for Wuhan ancestral with the D614G mutation and D variant were performed as previously described (19). Spike protein from different variants were purified using a one-step affinity method with NGL COVID-19 Spike Protein Affinity Resin (Repligen, Waltham, MA, USA) (19). The identity and purity of the antigens was also confirmed by mass spectrometry. Absence of endotoxin contamination was verified using Endosafe cartridge-based Limulus amebocyte lysate tests (Charles River Laboratories, Charleston, SC, USA).

### Mice

In SARS-CoV-2 challenge studies, the K18-hACE2 transgenic mice were used. Mice were obtained from Comparative Medicine Centre (CONICET -National University of the Litoral) and housed in a BSL-3 Laboratory Animal Facility (ANLIS, Dr. Carlos Malbrán, Buenos Aires) under a 12 h light/dark cycle, with free access to food and water. All procedures were performed in accordance with protocols approved by the Ethical Committee for Animal Experiments of the Faculty of Science at La Plata National University (Argentina, approval number 010-38-21 y 004-40-22).

### Vaccination and SARS-CoV-2 Challenge

Spike proteins (2 μg/dose) were formulated with Alhydrogel 2% (InvivoGen, San Diego, CA, USA) as previously described (20). Vials of AZD1222/ChAdOx1 (Astra Zéneca, AZD1222), Spikevax/mRNA-1273 (Moderna, mRNA-1273) were acquired from Ministry of Health of Buenos Aires province. Female and male mice were randomly divided into immunized and non-immunized groups with 6 to 8 mice in each group. Briefly, a primary immunization scheme consisting of two doses of a commercial vaccine with an interval of 14 days, followed by a third booster dose applied 2 months after the second dose. For the primary schedule, mice were intramuscularly immunized with of AZD1222 using 1/10 of the Human Dose (HD). For the booster, commercial vaccine or 2 μg/dose of ancestral W or D Spike proteins were used. Two weeks post boost, mice were anesthetized with inhalation isoflurane and challenged intranasally (i.n.) with 5x10^5^ – 5x10^6^ PFU of SARS-CoV-2 strain Omicron BA.1 (25 μl per nare). Animals were weighed and monitored daily for any clinical signs of disease. On day 5 post infection, animals were euthanized to examine viral replication in the respiratory tract. Brain and lungs were also collected for histopathologic studies. Euthanasia was performed by cervical dislocation. The procedures were carried out in BSL-III facilities.

### Sample collection and tissue harvest

For the assessment of humoral immune responses, blood was collected from isoflurane-anesthetized mice via the submandibular vein at different time after a vaccination dose. At sacrifice, blood, spleen, brain, nasal wash, and lungs were collected. Blood was centrifuged at 8,000 g for 10 min to separate serum. The brain and the left lung were collected for histopathological examination, while the right lung and the nasal wash were collected to determine the viral titter using RT-qPCR assay. Spleens were processed into single-cell suspensions, washed, and resuspended in RPMI medium supplemented with 1% penicillin/streptomycin and 10% FBS to determine cellular response and perform cytometric analyses. The procedures were carried out in BSL-III facilities.

### Histopathological analysis

Tissues of challenged K18-hACE-2 mice were fixed in a buffer containing 4% formalin. Sections were prepared and stained with hematoxylin and eosin (H&E). Lung sections were scored on a severity scale of mild, moderate, and severe based on the degree of inflammatory cell infiltration at various anatomical sites, which could be focal or multifocal. Lung disease at different anatomical sites was assessed under light microscopy by evaluating the expansion of the parenchymal wall and the presence of interstitial pneumonia, alveolitis, and bronchiolitis. The severity of brain tissue lesions was determined by their extent and scored on a severity scale of mild, moderate, or severe based on the degree of inflammatory cell infiltration at different anatomical sites in the brain. Specifically, the presence of white matter vacuolization, dark neurons, haemorrhage, and congestion were analysed.

### Anti-spike IgG ELISA

Anti-spike total IgG titers and IgG isotypes in serum were quantified by ELISA. Briefly, 96-well high-binding ELISA plates (Nunc A/S, Roskilde, Denmark) were coated overnight at 4ºC with 100 μL of 0.45 μg/mL Spike protein. Plates were washed five times with PBS/0.05% Tween20 (PBS-T), and then blocked for 1 h at 37 °C with 200 μL 3% milk in PBS before incubation with serially diluted samples of mouse serum (1 h, 37°C). After five washes with PBS-T (Sigma-Aldrich), 100 μL of horseradish-peroxidase–labelled goat anti-mouse IgG (Invitrogen, United States) at 1:8,000. For measuring the IgG isotypes, the bound antibody was incubated with horseradish-peroxidase labelled subclass-specific anti-mouse IgG1 at 1:8,000 or anti-mouse IgG2c at 1:2,000 (Sigma, Aldrich). After five washes with PBS-T, 100 μL/well of the substrate o-phenylenediamine dihydrochloride (OPD, Sigma-Aldrich) diluted in 0.05 M citrate buffer (pH 5.0) was added. Plates were developed for 15 min at RT in the dark. The reaction was stopped with 50 μL/well of 4 N H_2_SO_4_. Absorbances were measured with Titertek Multiskan Model 340 microplate reader (ICN, USA) at 490 nm. From the experimental protocol performed in triplicate, one representative experiment is presented in the Results.

### Pseudovirus neutralization assay

Neutralization assays were carried out with SARS-CoV-2 pseudotyped particles (CoV2pp-GFP from Sean Whelan laboratory) (21) that carries vesicular stomatitis virus as viral backbone and expresses full length wild-type spike from Wuhan ancestral or Omicron variant on its envelope. Viral stocks (VSV-eGFP-SARS-CoV-2) were amplified using 293T ACE2/TMPRSS2 cells at an MOI of 0.01 in Dulbecco’s Modified Eagle’s medium containing 2% FBS at 37ºC. For neutralization assays, Vero cells maintained with DMEM high glucose with 10% FBS were seeded in a 96-well plate the day before infection. Serum neutralizations were performed by first diluting the inactivated sample 2-folds and continuing with a 2-fold serial dilution. A pre-titrated amount of pseudotyped particles was incubated with a 2-fold serial dilution of patient sera for 1 h at 37 ºC prior to infection. Subsequently, cells were fixed in 4% formaldehyde containing 2 mg/mL DAPI nuclear stain (Invitrogen) for 1 hour at room temperature, and fixative was replaced with PBS. Images were acquired with the InCell 2000 Analyzer (GE Healthcare) automated microscope and analysed as previously described (20). Absolute inhibitory concentrations (absIC) values were calculated for all animals’ sera samples by modelling a 4-parameter logistic (4PL) regression with GraphPad Prism 9. Absolute inhibitory concentration was calculated as the corresponding point between the 0% and 100% assay controls. Fifty % inhibition was defined by the controls for all the samples on the same plate. The procedures were carried out in BSL-II facilities.

### Ag-specific IFN-γ, IL-17 and IL-5 production by spleen cells and FACS analysis

Spleens from untreated and immunized mice were passed through a 40-mm cell strainer to obtain a single-cell suspension. Spleen cells were cultured with W, and D spike proteins (1.5 μg/ml), or medium only. After 72 h of incubation, IFN-γ, IL-5 and IL-17A concentrations were quantified in supernatants by ELISA.

For FACS analysis, spleen cells that were not re-stimulated with Spike protein were incubated with CD16/CD32 FcgRIII (1:100) to block IgG Fc receptors. Cells were treated with LIVE/DEAD Violet (Invitrogen), followed by surface staining with fluorochrome-conjugated anti-mouse Abs for various markers: CD3-Pe-Cy7 (BD), CD8-PE (BD), CD4-FITC (BD), CD44-PE (BD), CXCR5-PerCP-Cy5.5 (BD), CD127-APC (BD). Flow cytometry analysis was performed on an BD FACSAria Fusion. The results were analyzed using FlowJo software (TreeStar). The procedures were carried out in BSL-II facilities.

### Detection of SARS-CoV-2 in tissues

For detection of viral RNA, tissues collected were stored in RNAlater. RNA was extracted using a QIAamp Viral RNA Mini Kit (Qiagen, Hilden, Germany), according to manufacturer’s instructions. RT-qPCR detection of SARS-CoV-2 was performed using kit LightMix® Modular SARS-CoV-2 (COVID19) E-gene, LightMix® Modular SARS-CoV (COVID19) RdRp (TIB MOLBIOL), and RNAsa P following the protocol developed by Dr. Corman et al from the Institute of Virology, Charité, Berlin, Germany, by the World Health Organization (https://www.who.int/docs/default-source/coronaviruse/protocol-v2-1.pdf) on a real time thermal cycler CFX96 (Biorad).

The measurement of the viral loads for different SARS-CoV-2 genes in tissue homogenates was be carried out as previously described (22). The results were normalized considering the exact weight of the organ or volume of the fluid.

### Statistical analysis

All statistics were analyzed using GraphPad Prism 9. Ordinary one-way ANOVA was performed to determine statistical significance between groups of three or more (Tukey’s multiple comparison test or Bonferroni’s multiple comparison test as appropriate). For fluorescence values analysis, we used Mann-Whitney statistical analysis. Statistical differences between groups of two were determined via unpaired t test for data deemed normal. Differences were significant when p< 0.05.

## Results

We have previously demonstrated the immunogenicity capacity of the SW-Vac 2μg vaccine candidate in a mouse model used for primary immunization schemes as well as boosters (20). This vaccine contains the soluble glycosylated trimeric prefusion stabilized spike protein derived from the ancestral virus. Taking into account that Omicron sub-lineages can evade polyclonal neutralizing antibody responses induced by the primary vaccine series, but boosters have demonstrated effectiveness against severe disease caused by Omicron, we decided to evaluate the protective capacity of our vaccine candidate when used as a heterologous booster dose (66, 67). Vaccination schemes that included 2 doses spaced 14 days apart and a third booster dose applied 2 months after the second dose were used (Fig 1A). For the primary series, 2-dose regimens of mRNA1273 or AZD1222 were used. As a comparative measure, SD-Vac 2μg (containing trimeric S protein for Delta variant) was also employed as a booster dose. The experiments were conducted in the murine model using transgenic mice, as wild-type laboratory mice are not highly susceptible to SARS-CoV-2 infection due to the mouse angiotensin-converting enzyme (ACE) 2 enzyme not acting as a cellular receptor for the virus (23). Several transgenic mouse strains expressing the human version of the SARS-CoV-2 receptor (hACE2) replicate viral replication and certain clinical characteristics of human infection (23). The most commonly used model, which we also employed, is the K18-hACE2 transgenic mouse (24), which predominantly expresses hACE2 in epithelial cells under the control of the cytokeratin 18 (KRT18) promoter (25).

**Figure 1.**
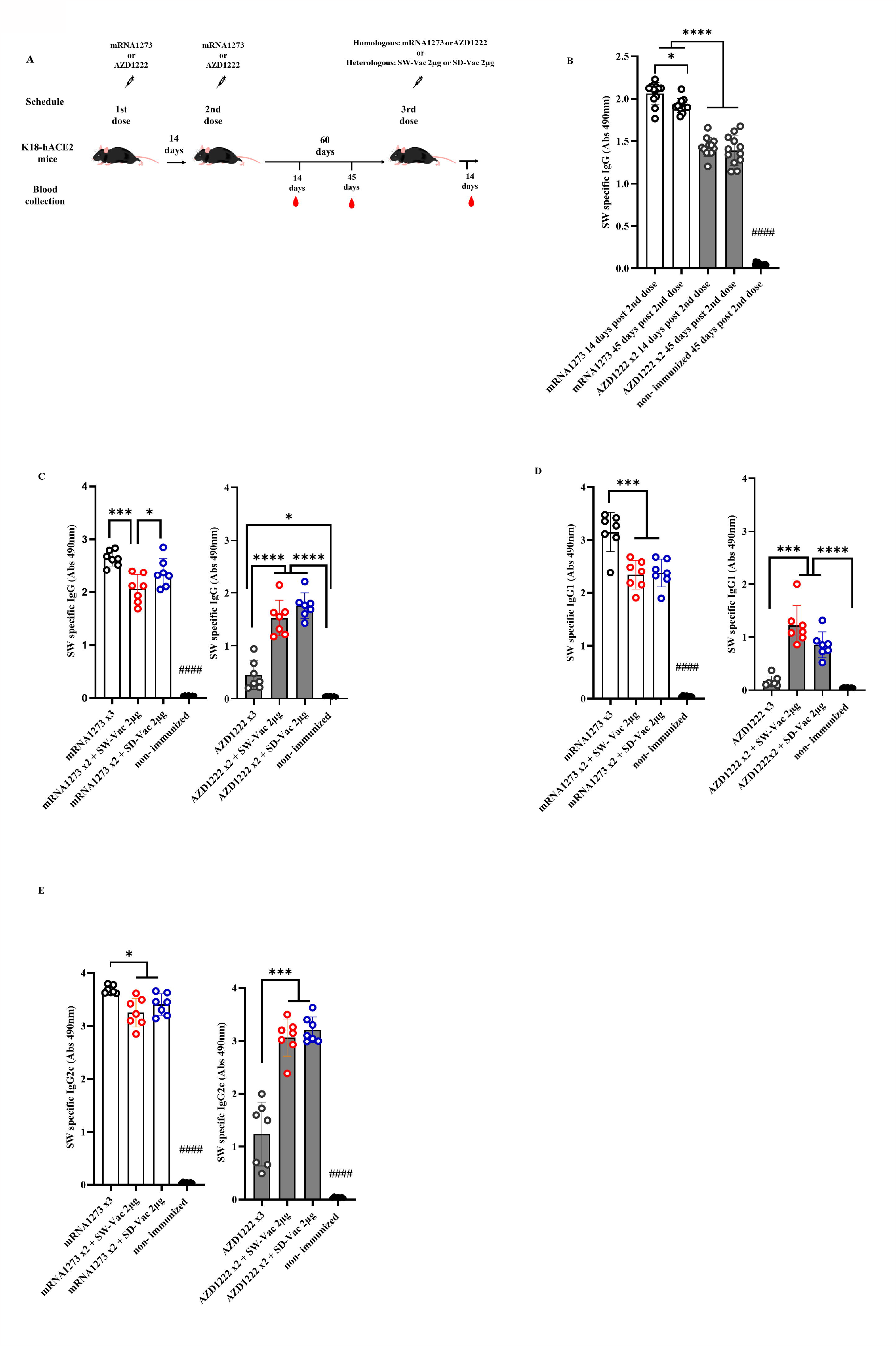
Specific ancestral Spike (SW) IgG and IgG isotypes levels induced by homologous and heterologous vaccination schedule. **A)** K18-hACE2 mice (n =7/group) were immunized on Days 0 and 14 with commercial vaccines mRNA1273 or AZD1222 plus a booster with formulations containing purified spike protein from ancestral (SW) or Delta (SD) SARS-CoV-2 variant on Day 60 delivered intramuscularly. **B)** SW specific IgG levels induced by 2 dose primary series. Serum collected on Days 14 and 45 after the second dose was analyzed by ELISA. **C)** SW specific IgG, **D)** IgG1 and **E)** IgG2c levels induced by 3-dose homologous and heterologous schemes. Sera from immunized and non-immunized were collected 14 Days after the 3^rd^ dose and analyzed by ELISA. Serum antibodies levels are expressed as the geometric mean of Abs490 nm ± SD from each group. For statistical analysis, antibody levels were analyzed by a one-way ANOVA with Bonferroni’s multiple comparisons test. The * symbol indicate significant differences among immunized: ****p<0.0001, *** p<0.001, *p<0.05. The # symbol indicate significant differences between non-immunized and immunized animals: #### p <0,0001.

To conduct a comparative analysis of the two primary vaccination series employed in this study, levels of SW-specific IgG were assessed at 14 and 45 days following the second dose, using ELISA assays. The results are presented in Fig 1B. While IgG levels for the non-immunized mice group were undetectable, significantly high IgG levels were observed for all immunized groups, with the two-dose mRNA1273 treatment displaying the highest levels (p<0.001). Almost in all immunized groups, SW specific IgG levels detected at 14 days after the second immunization did not substantially differ from those at 45 days following the second dose. Only a small reduction of 7% was detected for the mRNA 2-dose scheme at 45 days post immunization in comparison with that obtained at 14 days post immunization (p<0.05). The levels of SW-specific IgG for the mRNA scheme were significantly higher than those induced by the AZD1222 scheme (Fig 1B). This difference between the vaccines tested here was also observed for the homologous 3-dose schemes (Fig 1C). For the schemes with two doses of mRNA1273 + a SW-Vac 2μg booster, the levels of SW-specific IgG were slightly lower than those from animals treated with 3 doses of mRNA1273 (p<0.001) (Fig 1C). As previously described (20), in the case of the non-replicative viral vector vaccine AZD1222, the levels of SW-specific IgG detected in the sera of mice immunized with heterologous regimens (AZD1222 x2 + SW-Vac 2μg or AZD1222 x2 + SD-Vac 2μg) were significantly higher than those detected in mice treated with the homologous 3-dose AZD1222 regimen (p<0.0001) (Fig 1C). The levels of IgG induced by the homologous or heterologous AZD1222 schemes were lower than those detected in mRNA1273 vaccinated mice (p<0.001).

The homologous 3-dose mRNA1273 scheme also induced the highest levels of SW-specific IgG1 (Fig 1D). The heterologous schemes including mRNA1273 were slightly lower than those induced by the homologous 3-dose mRNA1273 regimen (p<0.001 for mRNA1273 x2 + SW-Vac 2μg vs. mRNA1273 x3; p<0.01 for mRNA1273 x2 + SD-Vac 2μg vs. mRNA1273 x3). In contrast, for the homologous scheme with 3 doses of the commercial AZD1222 vaccine, the levels of SW-specific IgG1 were the lowest (Fig 1D). The heterologous scheme including AZD1222 + SW-Vac 2μg induced levels of SW specific IgG1 that were significantly higher than those of the homologous 3-dose AZD1222 regimen. Regarding the levels of SW-specific IgG2c, the highest levels were observed in the treatment groups that received doses of mRNA1273, both in homologous and heterologous regimens, as well as in the heterologous treatment groups based on AZD1222 (Fig 1E). The lowest levels were detected for the homologous 3-dose regimen with AZD1222 (Fig 1E). Similar results to those mentioned above regarding IgG, IgG1, and IgG2c levels were obtained when the evaluations were conducted against SD (refer to supplementary material, Fig 1S).

When comparison between IgG2c levels with respect to IgG1 were performed, an interesting finding was that for all evaluated homologous and heterologous schemes the IgG2c/IgG1 ratios were higher than 1 (supplementary material Table S1), suggesting the induction of a Th1-directed immune response.

Neutralizing capacity was also evaluated by measuring antibody-neutralizing titers against W ancestral variant, using the CoV2pp-GFP pseudovirus infection (Fig 2 A-D). The highest neutralizing antibody titers were observed in the 3-dose mRNA1273 scheme. The heterologous schemes (3 doses) using either mRNA1273 or AZD1222 induced higher levels of neutralizing antibodies compared to the homologous 3-dose AZD1222 scheme. Specifically, for the AZD1222 x2 + SW-Vac 2μg heterologous scheme, the neutralizing antibody titers were approximately 9 fold higher than those detected in mice immunized with the AZD1222 x3 scheme (3218.29 ± 413.8 vs 365.7 ± 51.72, p<0.0001), and 1.37 fold higher than those observed for the AZD1222 x2 + SD-Vac 2μg scheme (p<0.05). The lowest levels of neutralizing antibodies were detected in the groups treated with AZD1222 in 2 or 3 doses (Fig 2 B). In the non-immunized group of animals, no detectable titers of neutralizing antibodies were found. These assays were repeated using the pseudotyped virus expressing the S protein of the Omicron (BA.5) variant of the SARS-CoV-2 virus. The results presented in Fig 2C and D show that the treatments incorporating mRNA1273 doses, whether in homologous 2- or 3-dose or heterologous 3-dose schemes, exhibited significantly higher antibody titers compared to the 2-dose primary regimen involving the mRNA1273. Conversely, the AZD1222 x 2-dose and 3-dose schemes displayed lower antibody levels than the heterologous scheme either with SW-Vac 2μg or SD-Vac 2μg (p<0.001, Fig 2D). In the group of non-immunized animals, neutralizing antibodies were not detected.

**Figure 2.**
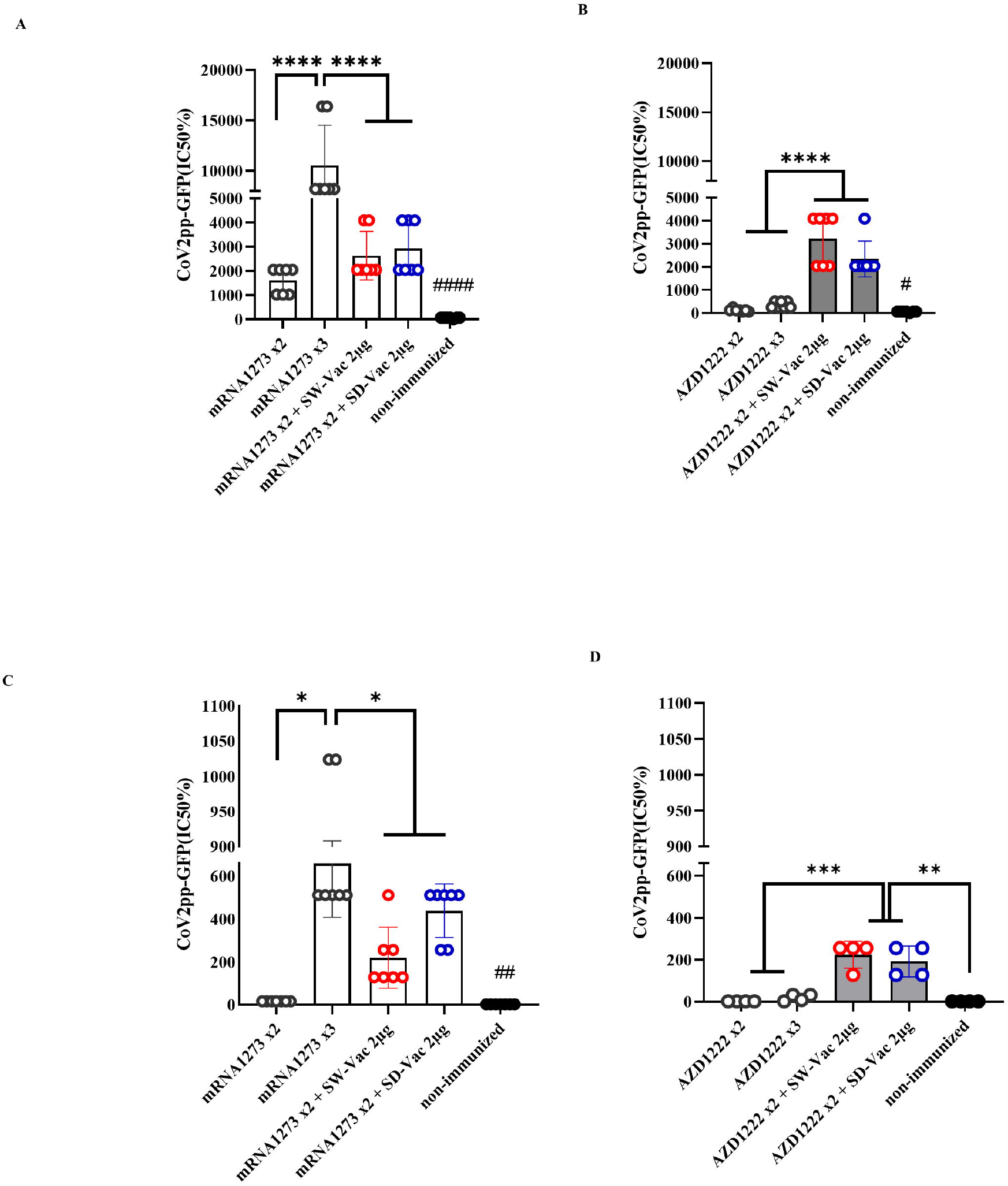
Neutralizing antibodies induced by homologous and heterologous vaccination schedule. Mice were immunized as shown in figure 1 panel A. Serum was collected on Day 14 after the 2^nd^ and 3^rd^ dose. Neutralizing titers were measured by 50% inhibition for the pseudotyped virus (CoV2pp-GFP) expressing W **(A** and **B)** or Omicron **(C** and **D)** spike protein. Data from mRNA1273 vaccinated mice are presented in panels **A** y **C** and those from AZD1222 in panels **B** and **D**. Absolute inhibitory concentration was calculated as the corresponding point between the 0% and 100% assay controls. Fifty % inhibition was defined by the controls for all the samples on the same plate. For statistical analysis, values were analyzed by a one-way ANOVA with Bonferroni’s multiple comparisons test. The * symbol indicate significant differences among immunized: **** p<0.0001, ***p<0.001, **p<0,01, * p<0.05. The # symbol indicate significant differences between non-immunized and immunized animals: #### p<0.0001, ##p<0,01, #p<0,05,

The groups of animals that received 2-dose regimens plus a booster were intranasally challenged with a suspension of SARS-CoV-2 (5x10^5^ – 5x10^6^ PFU Omicron BA.1/40 μl) 14 days after receiving the booster dose (Fig 3A). Five days post-challenge, the mice, including those in the non-immunized group, were euthanized by cervical dislocation to evaluate the cellular immune response and colonization (viral load) in the respiratory tract. Additional samples were taken for histological evaluations.

**Figure 3.**
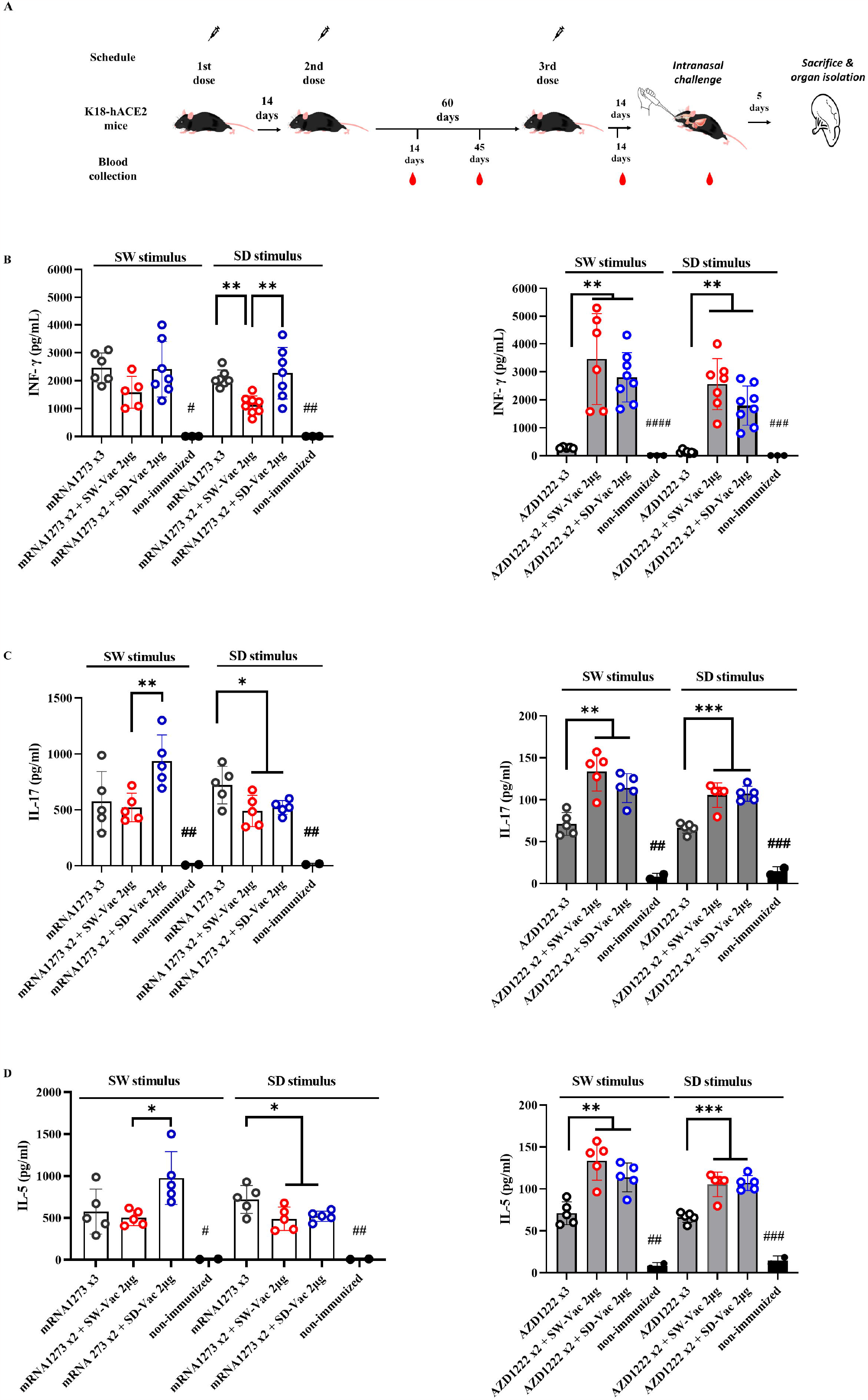
Homologus and heterologous immunization induces Ag-specific Th1 (IFN-γ) / Th17 (IL-17) / Th2 (IL-5). **A)** K18-hACE2 mice (n =7/group) were immunized on Days 0 and 14 with commercial vaccines mRNA1273 or AZD1222 plus a booster with formulations containing purified spike protein from ancestral (SW) or Delta (SD) SARS-CoV-2 variant on Day 60 delivered intramuscularly. Mice were challenged 14 days after the 3^rd^ dose with SARS-CoV-2 Omicron BA.1 variant, 5 days after challenge mice were sacrificed and spleens were harvested. Levels of secreted IFN-γ **B)**, IL-17 **C)** and IL-5 **D)** following splenocytes stimulation with medium or recombinant spike protein from W o D variants were determined by ELISA. Bars are means ± SEM of pg/ml. The * symbol indicate significant differences among immunized: *** p<0.001, **p<0.01, *p<0.05. The # symbol indicate significant differences between non-immunized and immunized animals: #### p<0.0001, ### p<0.001, ## p<0.01, # p<0.05

Regarding the cellular immune response, we conducted splenocytes proliferation assays using SW and SD as stimuli at a final concentration of 1.5 μg/ml. After the incubations with the stimuli and in the non-stimulated controls, the supernatants were collected to evaluate the secreted levels of IFN-γ (Th1 profile marker), IL-5 (Th2 profile marker), and IL-17 (Th17 profile marker) using ELISA assays (Fig 3 B-D). The vaccination schedules that included doses of mRNA1273, either in a homologous or heterologous manner, elicited similar levels of IFN-γ when stimulated with the S protein from the ancestral virus and higher than those observed in the non-immunized group (Fig 3B). On the other hand, when stimulated with the SD protein, a significant difference was observed in the heterologous scheme with the SW-Vac 2μg booster, which showed significantly lower levels compared to the mRNA1273x2+SD-Vac 2μg and mRNA1273 treatments (1585 ± 255.1 vs 2411 ± 375.0 and 2458 ± 219.1, respectively; p<0.01). Regarding IL-17 and IL-5, when analyzing the data in response to SW and SD stimuli, it was observed that all immunized animals had significantly higher levels than those detected in the non-immunized control group (Fig 3C and 3D). For both interleukins, it was found that, in the case of the SW stimulus, levels in the heterologous scheme with SD-Vac 2μg booster were higher than those in the heterologous scheme with SW-Vac 2μg booster, which, in turn, did not show differences with the homologous scheme of 3 doses of mRNA1273. For the SD stimulus, the heterologous schemes yielded slightly lower levels of IL-17 (Fig 3C) and IL-5 (Fig 3D) compared to those detected in the homologous scheme (p<0.05).

In the case of treatments that included AZD1222, it was observed that in heterologous schemes, levels of IFN-γ were significantly higher compared to the homologous scheme of 3 doses, both against SW and SD stimulus (p<0.01, Fig 3B). The levels of IFN-γ in the homologous scheme, although low, showed a significant difference compared to the non-immunized group (Fig 3B). For AZD1222 schemes a similar behavior was observed in the levels of IL-17 and IL-5 (p<0.01). For both, SW and SD stimulus heterologous schemes that included AZD1222 induced significantly higher levels of both IL-17 (Fig 3C) and IL-5 (Fig 3D) compared to those detected for the homologous scheme. For these interleukins, the levels detected in the homologous scheme were higher than those in the non-immunized control group (p<0.05). It is worth noting that, for both interleukins, the levels detected in all schemes that included AZD1222, whether heterologous or homologous, were significantly lower than the levels detected in regimens that included mRNA1273.

All these results demonstrate that the tested immunization schedules induced a mixed Th1/Th17/Th2 profile. In the schedules using AZD1222 as the primary doses, higher levels of interleukins were observed in the heterologous schedules compared to the homologous schedule. Furthermore, the heterologous schedules, except for the schedule with SD-Vac 2μg with SD stimulus, exhibited higher levels of Th1 marker (IFN-γ) than those detected in the schedules using mRNA1273, while IL-5 and IL-17 levels were higher in the schedules including mRNA1273 in compassion with AZD1222 schemes.

We also evaluated the population of CD4 Teff, CD4 Tcm and CD4 Tfh in the spleens of immunized and non-immunized challenged animals (Fig 4). The CD4 T flow cytometry analysis is presented in supplemental material Fig 2S. We found that all treatments induced significantly higher levels of CD4 Teff compared to the non-immunized control group (p<0.05, Fig 4A). Furthermore, within the homologous scheme, the highest levels were observed for the mRNA1273x3 treatment, which are significantly different from those detected for AZD1222x3 (p<0.01). While for the mRNA1273-based schemes, the levels of this population were indistinguishable between the heterologous and homologous treatments (Fig 4A), for the AZD1222-based treatments, the heterologous schemes with SW-Vac 2μg yielded significantly induced higher CD4 Teff levels compared to the corresponding AZD1222x3 homologous (p<0.05). Regarding the CD4 Tcm population, we observed that all treatments induced significantly higher levels of this population compared to the non-immunized control group (p<0.05, Fig 4B). No differences were observed between the homologous or heterologous schemes of the two vaccine platforms tested (Fig 4B). All treatments also significantly induced higher CD4 Tfh population compared to the control group (p<0.05). For the schemes that included doses of mRNA1273 or AZD1222, no differences in CD4 Tfh population were detected between the homologous and heterologous schemes (Fig 4C).

**Figure 4.**
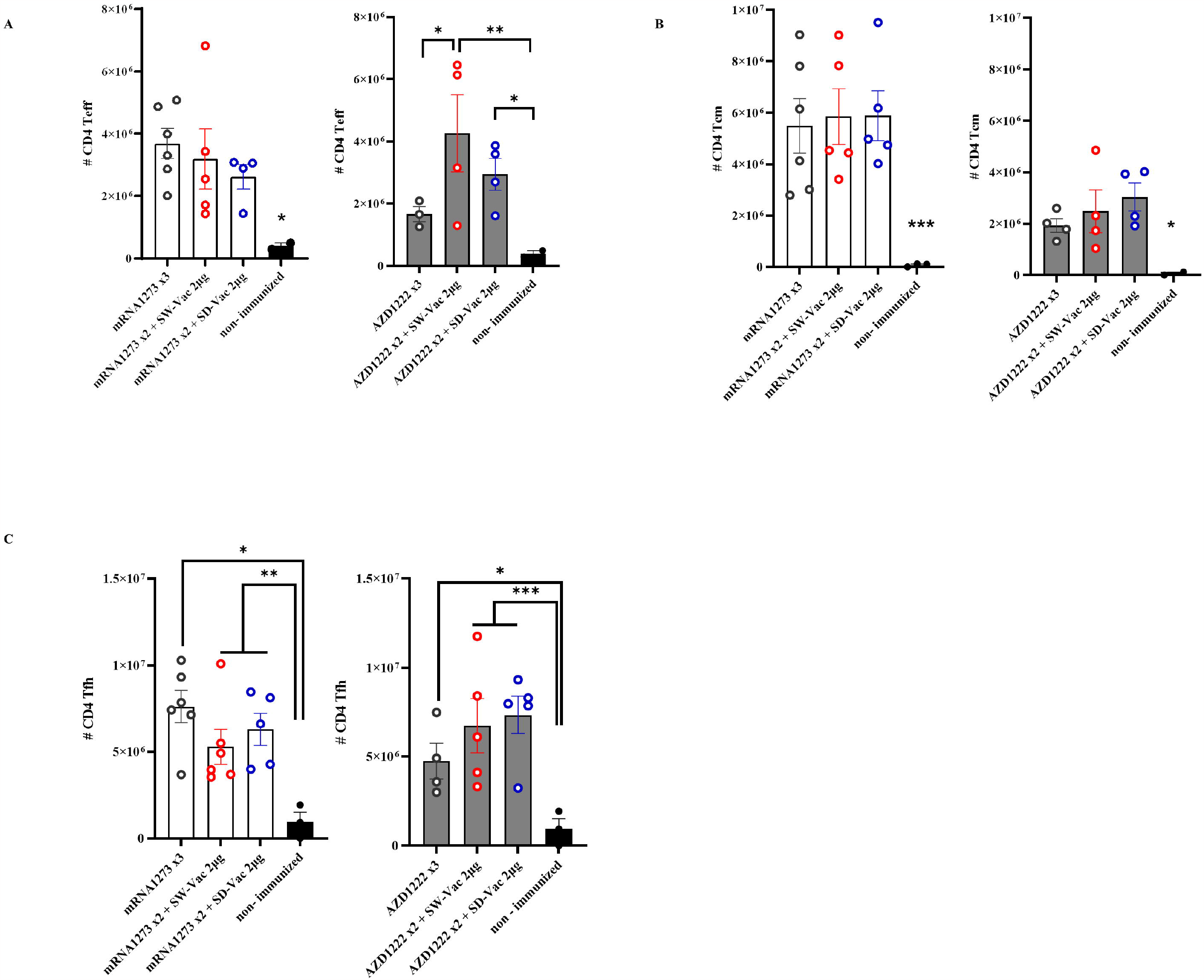
T CD4+ populations induced after Omicron BA.1 challenge in mice vaccinated with homologous or heterologous schemes. Five days post challenge immunized and non-immunized mice were sacrificed and spleen cells were collected. For FACS analysis the spleen cells were incubated with CD16/CD32 Fc g RII to block IgG Fc receptors. Cells were incubated with LIVE/DEAD Violet (Invitrogen), followed by surface staining with fluorochrome-conjugated anti-mouse Abs for various markers. **A)** CD4+CD44+CD62L-, (CD4+ Teff), **B)** CD4+CD44+CD62L+ (CD4+ Tcm) and **C)** CD4+CXCR5+(CD4+ Tfh). Flow cytometry analysis was performed on an BD FACS Aria Fusion. The results were analyzed using Flow Josoftware (TreeStar). The represented data correspond to the mean of absolut cells counts ± SEM. Two-way ANOVA followed by Bonferroni post-test were used for statistical analysis. ***p<0.001; **p < 0.01, *p < 0.05.

Throughout the entire assay, the weight of mice belonging to different groups was monitored (Fig 5B and E). All mice showed increased body weight until day 75 of treatment initiation, with weights remaining constant thereafter. On day 90, after challenge with a suspension of SARS-CoV-2 Omicron BA.1 (2x105 PFU Omicron BA.1/40 μl) in non-immunized animals, a continuous decrease in weight was observed, with a drop of approximately 3% at 4 days post-challenge (4dpc). In the schemes using mRN1273 in the primary series, a weight drop was detected for the group whose regimen included SD-Vac 2μg as a booster, which was quite similar to the non-immunized group. No weight loss was detected in the homologous 3-dose schemes, and a slight increase was observed in the heterologous scheme with SW-Vac 2μg, although it did not differ from the homologous 3-dose treatment. In the case of the groups immunized with a homologous or heterologous booster that included AZD1222 as the primary series, a continuous reduction in weight similar to that observed in the non-immunized infected group was detected (approximately 2% on day 4 post challenge).

**Figure 5.**
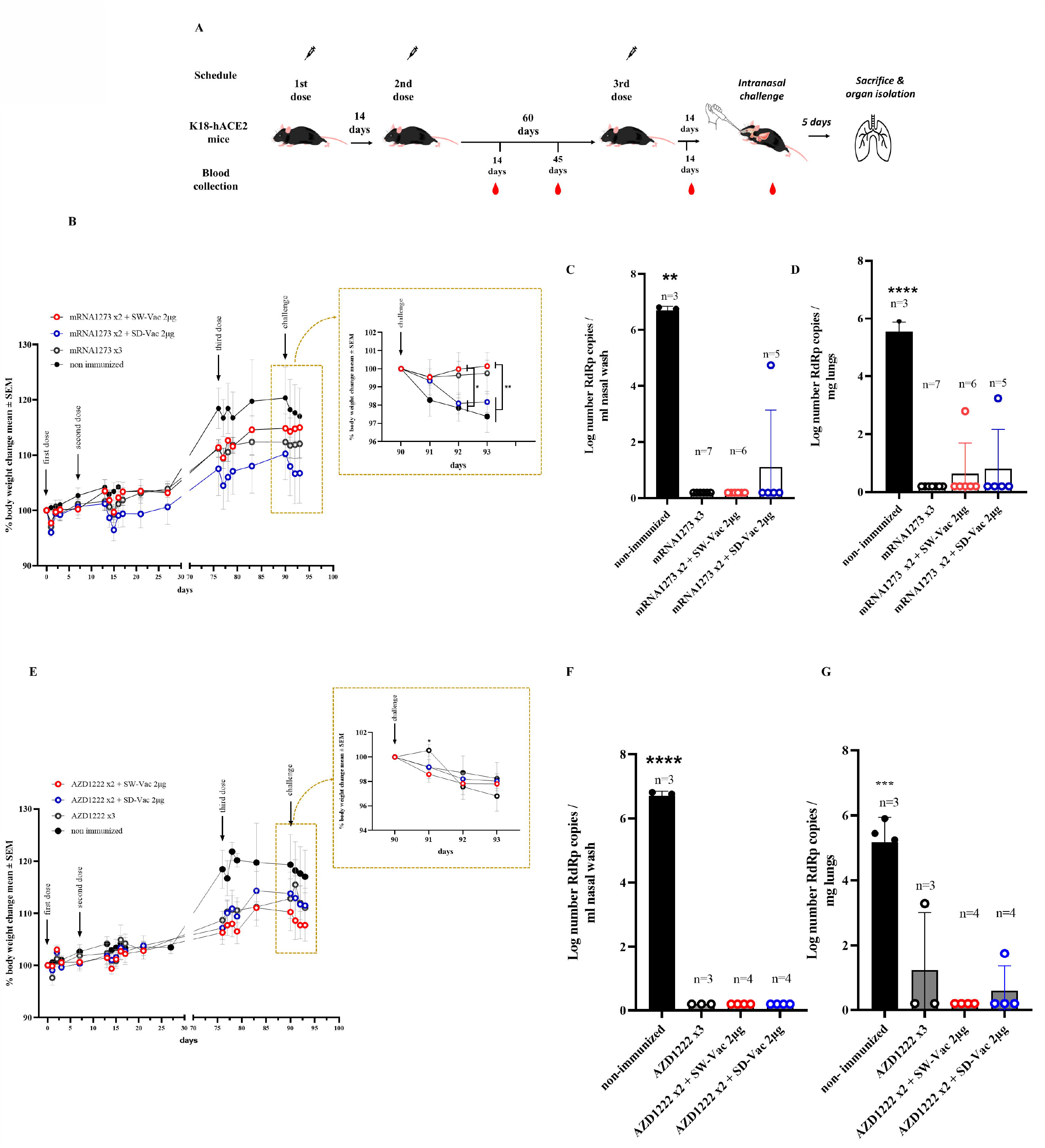
Protection assay against Omicron BA.1 SARS-CoV-2 infection in the intranasal challenge mouse model. **A)** K18-hACE2 mice (n =7/group) were immunized on Days 0 and 14 with commercial vaccines mRNA1273 or AZD1222 plus a booster with formulations containing purified spike protein from ancestral (SW) or Delta (SD) SARS-CoV-2 variant on Day 60 delivered intramuscularly. Mice were challenged 14 days after the 3^rd^ dose with SARS-CoV-2 Omicron BA.1 variant, 5 days after challenge mice were sacrificed and nasal washes and lungs were collected. **B and E)** Changes in body weight were expressed as the percentage of body weight in comparison to day 0 (mean ± SEM.). Post-challenge weights were zoomed in for each treatment in the right part of the graph. **C** and **F)** nasal and **D** and **G)** lungs SARS-CoV-2 viral load after Omicron BA.1 challenge in animals vaccinated with schedules that contain mRNA-1273 (C and D) or AZD1222 (F and G). Results are expressed as Log number Rdrp copies/ml nasal wash or as Log number Rdrp copies/ mg lungs. One-way or Two-way ANOVA followed by Bonferroni post-test, were used for statistical analysis. ****p < 0.0001, ***p<0.001, **p < 0.01.

In terms of virus load, at 5 dpc, mice in control group had a high level of 4.783 10^6^ copies/mL in nasal wash (Fig 5 C and F) and 4.21 x10^5^ copies/mg in the lungs (Fig. 5 D and G). The majority of immunized mice (28/29) showed a significant reduction (almost 7 orders of magnitude, p<0.001) in viral load in the nasal wash compared to non-immunized animals. Only one immunized mouse, which received the mRNA1273x2 + SD-Vac 2μg regimen, exhibited a viral load of 4.74 x 10^4 copies/mL in the nasal wash (Fig 5C). Regarding viral load in the lungs, out of all the immunized animals (n=29), 25 mice showed a reduction of nearly 6 orders of magnitude in viral load compared to the control animals (Fig 5 D and G). Only 4 mice achieved a reduction of approximately 2.5 orders of magnitude. In 2 mice that received the primary regimen mRNA1273 + SW-Vac 2μg or SD-Vac 2μg, viral loads of 619 copies/mg and 1740 copies/mg, respectively, were detected in the lung tissue (Fig 5D). In another 2 mice that received a homologous regimen of 3 doses of AZD1222 or a primary regimen with that vaccine + 1 dose of SD-Vac 2μg booster, viral loads of 1930 copies/mg and 56 copies/mg of SARS-CoV-2 mRNA were detected in the lung tissue, respectively (Fig 5G).

Histological analysis was conducted at 5 days post-challenge. The non-immunized control group exhibited multifocal hyperplasia of type II pneumocytes, bronchiolar-alveolar hyperplasia, and infiltration of lymphocytes and monocytes in 100% of the animals’ lungs. In contrast, mice in the immunized group showed a reduction in pathological reactions and lung inflammation, indicating that the tested vaccination regimens may delay disease progression and result in milder lung inflammation during Omicron BA.1 infection. Among animals treated with vaccination regimens that included mRNA1273, only a maximum of 60% of the animals exhibited mild lung hyperplasia. Only 5.6% (1/18) and 11.1% (2/18) of the animals displayed moderate and multifocal hyperplasia of type II pneumocytes and bronchiolar-alveolar hyperplasia, respectively. In the case of animals immunized with regimens including AZD1222, 9.1% (1/11) exhibited hyperplasia of type II pneumocytes, and 18.2% (2/11) showed bronchiolar-alveolar hyperplasia. Furthermore, brain tissues from the control group presented moderate lesions, characterized by white matter vacuolization, dark neurons, and some samples displaying focal hemorrhage. However, immunized animals only showed significantly mild lesions. A moderate vacuolization of white matter was observed in only 38.9% (7/18) of animals immunized with mRNA1273-based schemes and in 63.6% (7/11) of those treated with AZD1222-containing schemes. No dark neurons or focal hemorrhages were detected in any case.

## Discussion

In this study, we evaluated the efficacy of a heterologous three-dose vaccination schedule in a mouse intranasal challenge model using the Omicron BA.1 variant of SARS-CoV-2. Two commercial vaccine platforms, mRNA1273 (mRNA-based vaccine) and AZD1222 (non-replicative vector-based vaccine AZD1222/ChadOx1), were used for a two-dose primary series. As a heterologous booster, we employed our vaccine candidate based on the glycosylated trimeric spike protein formulated with Alhydrogel (20). In a previous report, we demonstrated that the formulation containing the protein from the ancestral SARS-CoV-2 variant at a dose of 2 μg (SW-Vac 2μg), as well as the formulation containing the same dose of the Delta variant (SD-Vac 2μg), are immunogenic in both homologous and heterologous schemes, primarily when used as a heterologous booster in combination with the Gam-COVID-Vac-rAd26/rAd5 vaccine (20). Based on this, we used SW-Vac 2μg or SD-Vac 2μg as potential heterologous boosters against the Omicron BA.1 challenge. Homologous schemes of the two commercial vaccines included in this study were used as controls.

The evaluation of the immunogenic capacity in terms of the induction of specific IgG antibodies against the S protein from the ancestral or Delta variants of SARS-CoV-2 showed that the three-dose heterologous schemes proved to be comparable or more effective than the homologous schemes consisting in 3 doses of mRNA (SW-Vac 2μg 2.06±0.27 and SD-Vac 2μg 2.36±0.27 vs 2.64±0.14) or non-replicative vector platforms (SW-Vac 2μg: 1.52±0.33 and SD-Vac 2μg 1.76±0.24 vs 0.45±0.27), respectively (Figure 1). The differential IgG detected levels revealed their dependence on the vaccine platform used in the primary series. This distinct behavior was also observed in other markers of humoral immune response. The levels of IgG1 detected for the heterologous schemes on the primary AZD1222 scheme were higher than those detected for the corresponding homologous scheme. The heterologous boosters applied to the two-dose mRNA1273 primary series induced slightly lower levels of IgG1 compared to the corresponding homologous three-dose scheme. However, these levels were significantly higher than the homologous three-dose scheme of AZD1222, as well as the heterologous schemes with SW-Vac 2μg or SD-Vac 2μg that included the commercial formulation (p<0.001). An interesting finding was that for all evaluated homologous and heterologous schemes the IgG2c/IgG1 ratios were higher than 1 (supplementary material Table S1), suggesting the induction of a Th1-directed immune response. This Th1 profile was more pronounced in the case of the heterologous schemes that included the non-replicative vector vaccine AZD1222. These results are consistent with previous reports for these two vaccine platforms. In particular, in clinical trials and in *in vitro* studies using primary human PBMCs, MΦs, or moDCs, it has been observed that the AZD1222 vaccine induces much stronger interferon and pro-inflammatory cytokine gene expression than BNT162b2 and mRNA1273 mRNA vaccines (26–28). According to these results, in this study, we observed that the heterologous AZD1222 schemes elicited higher levels of INF-γ compared to the heterologous mRNA schemes after the Omicron BA.1 challenge (Figure 3). On the other hand, the homologous and heterologous schemes including mRNA vaccine doses, the levels of IL-17 (Th17 profile marker) and IL-5 (Th2 profile marker) were higher than those induced by the schemes including the non-replicative vector vaccine. Once again, the heterologous schemes including this commercial non-replicative vector platform induced significantly higher levels of IL-17 and IL-5 than the corresponding homologous 3-dose scheme. After the Omicron BA.1 challenge, it was observed that all vaccination schemes elicited a robust cellular immune response. This immune response involved the activation of various key components, including T follicular helper cells, effector memory T cells, and central memory T cells. These cellular immune responses are critical for the development of protective immunity (29).

It is worth noting that no significant differences were observed between SW-Vac 2μg and SD-Vac 2μg vaccines, demonstrating the cross-immunogenic reactivity capacity between the variants as we had previously detected (20).

The challenge with the Omicron BA.1 variant provoked in non-immunized mice a continuous weight loss, reaching a 3% loss at day 4 post challenge in accordance with the highest viral dose detected in nasal washes and lungs. This decrease was slightly lower in groups of mice immunized with a homologous or heterologous booster that included AZD1222 as the primary series. Furthermore, for the mRNA-based schemes, no weight loss was detected in the homologous 3-dose mRNA1273 schemes, and a slight increase was observed in the heterologous scheme with SW-Vac 2μg, although it did not differ from the homologous 3-dose treatment. Almost 97% of immunized mice (28/29) showed a significant reduction of 7 orders of magnitude in Omicron BA.1 viral load in the nasal washes compared to those detected in non-immunized animals (p<0.001). Moreover, more than 86% of the immunized animals (25/29) were completely protected against viral lung infection; only 4 mice achieved a reduction of approximately 2.5 orders of magnitude. The high protective capacity here observed are consistent with the neutralizing activity and the Th1 profile response detected, both of which have been reported as key factors in inducing protection (30). As expected the neutralizing capacity detected in the sera of immune animals was significantly higher against the pseudovirus expressing the ancestral variant S protein of SARS-CoV-2 than against the one expressing the S protein of the Omicron BA.1 variant (16 times less). Several studies also showed that Omicron was less sensitive to neutralization of immunized sera than D614G variant (31). Although the immune sera tested here exhibited reduced neutralization titers against the pseudovirus expressing the S protein of Omicron BA.1 variant, they were generally preserved to a significant extent. Furthermore, the third dose increased the Omicron neutralization titers. This phenomenon has been described by other authors, even in human populations, where it was observed that neutralization against the Omicron variant was lower compared to the wild type or Delta variant. However, it was generally found to be highest in individuals who recently received their third (booster) dose (32).

In the treatments that included AZD1222, the lowest levels of neutralizing antibodies against Omicron were observed, especially in the case of the homologous scheme. Nonetheless, the level of protection remained adequate and comparable to the other treatments that utilized the mRNA platform, likely due to the non-replicative vector vaccine inducing a higher Th1 profile. Based on the multiple protection markers induced by the schedules tested here, particularly those with mRNA primary series and the heterologous schedules with the AZD1222 vaccine, the vaccine’s efficacy could possibly be preserved over a longer period, even with a substantial reduction in neutralization of the immune sera.

In summary, the SW or SD trimer vaccine adjuvanted with Alhydrogel, which has been shown to be safe, demonstrated adequate humoral and cellular immune responses to induce protection against Omicron when administered as a booster. In the case of platforms like AZD1222, the heterologous booster with the vaccine prototypes presented here induce a more robust response than the homologous scheme that only includes AZD1222, potentially leading to an even higher long-term immunogenic difference. Thus, heterologous booster schemes prove to be a valid alternative for controlling variants such as Omicron and may even be superior compared to certain homologous schemes.

## Authors and Contributors

DFH planned the study, made the laboratory analysis, interpreted data, perform some experiments and drafted manuscript. AW, AG planned the study, interpreted data, and revised figures and the manuscript. MS purified the S proteins. YD expressed and purified the S proteins, revised figures and the manuscript. DB MEG, and MEZ performed certain experiments, interpreted data, and revised figures and the manuscript. PMA and ER performed experiments and laboratory analyses. MMGLL performed pseudovirus neutralization assays. All authors approved the final manuscript.

## Acknowledgments

We would like to acknowledge members of the Mammalian Cell Expression Section of the NRC-HHT for their contribution at producing, purifying and analyzing the spike proteins used in this study. Luciana Cayuela provided excellent technical assistance.

## Legends to the Figures

**Figure 1S.**
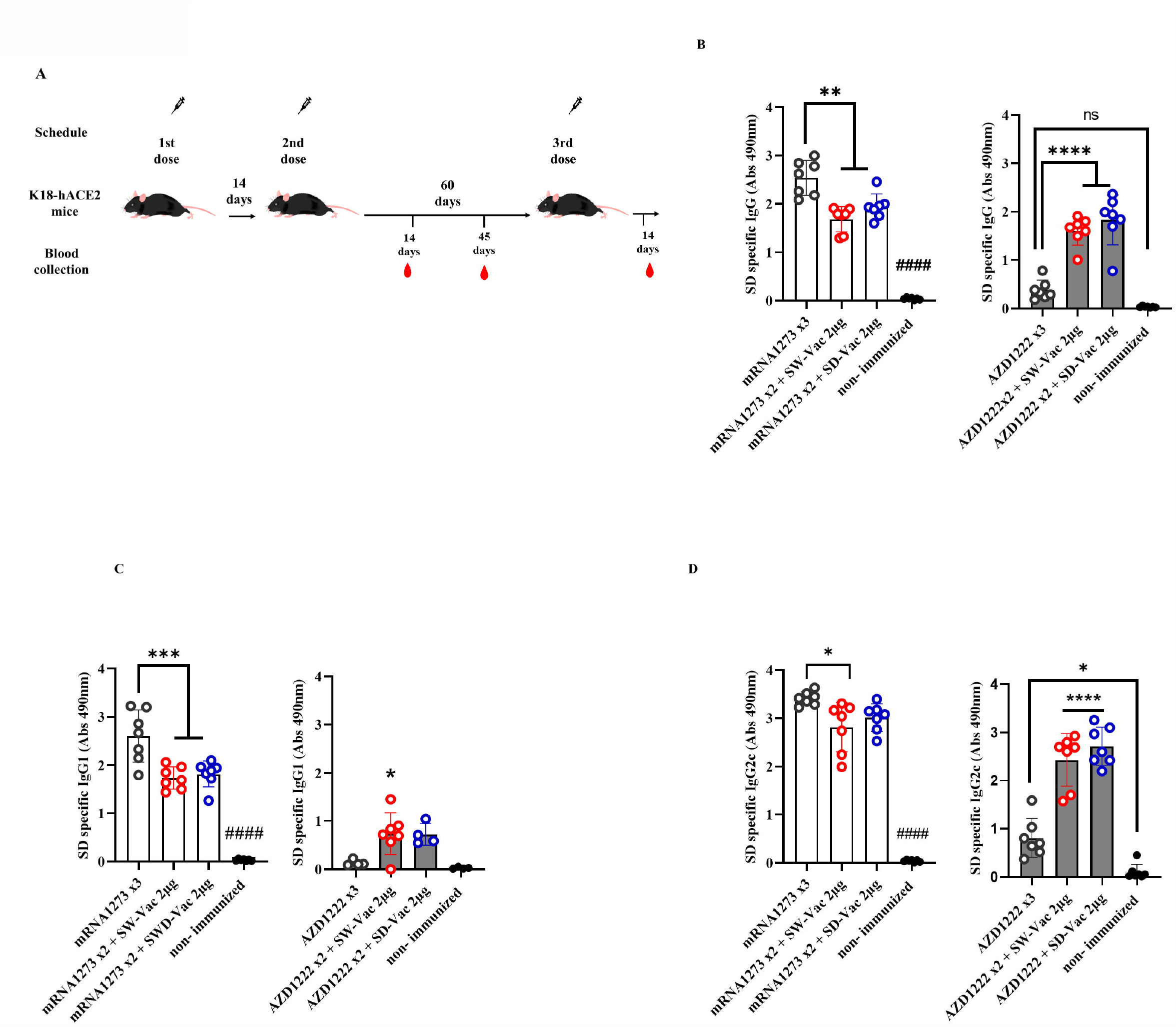
Specific ancestral Spike (SD) IgG and IgG isotypes levels induced by homologous and heterologous vaccination schedule. **A)** Mice vaccination schedule K18-hACE2 mice (n=7/group) were immunized on days 0 and 14 with anti SARS-CoV-2 commercial vaccines mRNA-1273 or AZD1222/ChAdOx1 and boostered at day 60 with formulations containing purified spike protein from ancestral (SW) or Delta (SD) SARS-CoV-2 variant delivered intramuscularly. SD specific **B)** IgG, **C)** IgG1 and **D)** IgG2c levels induced by 3-dose homologous and heterologous schemes. Sera from immunized and non-immunized were collected 14 days after the 3^rd^ dose and analyzed by ELISA. Serum antibodies levels are expressed as the geometric mean of Abs490 nm ± SD from each group. For statistical analysis, antibody levels analyzed by a one-way ANOVA with Bonferroni’s multiple comparisons test. The * symbol indicate significant differences among immunized: ****p<0.0001, *** p<0.001, *p<0.05. The # symbol indicate significant differences between non-immunized and immunized animals: #### p <0,0001.

**Figure 2S. CD4+ populations induced after Omicron BA.1 challenge in mice vaccinated with homologous or heterologous schemes**. Five days post challenge immunized and non-immunized mice were sacrificed and spleen cells were collected. Cells were incubated with LIVE/DEAD Violet (Invitrogen), followed by surface staining with fluorochrome-conjugated anti-mouse Abs for CD4+ T cells. A) Gate strategy for CD4+ population analysis by flow citometry. CD4+ effector T cells (CD3+CD4+CD44+CD62L-). CD4+ central memory T cells (CD3+CD4+CD44+CD62L+). CD4+ follicular helper T cells (CD3+CD4+CXCR5+). Flow cytometry analysis was performed on an BD FACS Aria Fusion. The results were analyzed using Flow Josoftware (TreeStar). B) Total number of CD4+ cells induced after Omicron BA.1 challenge in mice vaccinated with homologous or heterologous schemes. The represented data correspond to the mean of absolut cells counts ± SEM. Two-way ANOVA followed by Bonferroni post-test were used for statistical analysis. *p < 0.05.

**Table 1S.**
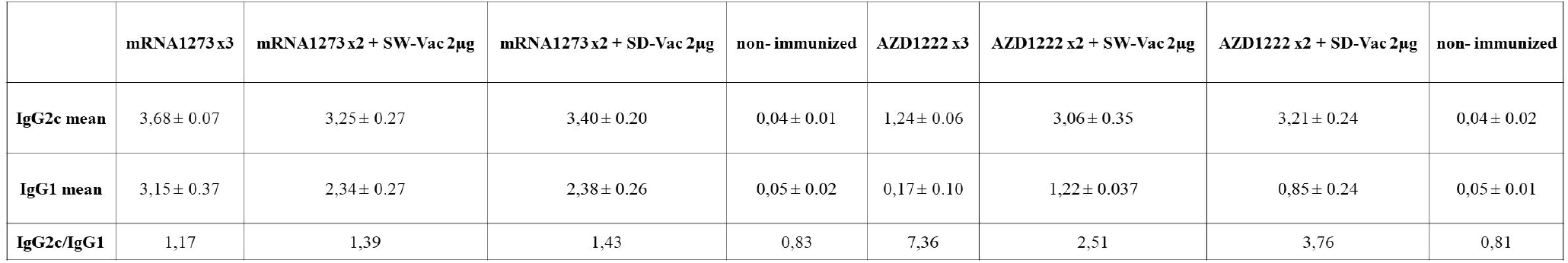
IgG2c and IgG1 titters and IgG2c/IgG1 ratio for homologous and heterologous immunized mice. Data for non-immunized mice are also included.

